# Temporal variation in the acoustic dynamic range is a confounding factor in EEG-based tracking of absolute auditory attention to speech

**DOI:** 10.1101/2025.03.04.641391

**Authors:** Tianyi Li, Simon Geirnaert, Davina Van den Broek, Elien Bellon, Bert De Smedt, Alexander Bertrand

## Abstract

Many studies have demonstrated that auditory attention to natural speech can be decoded from EEG data. However, most studies focus on *selective* auditory attention decoding (sAAD) with competing speakers, while the dynamics of *absolute* auditory attention decoding (aAAD) to a single target remains underexplored. The goal of aAAD is to measure the degree of attention to a single speaker, and it has applications for objective measurements of attention in psychological and educational contexts. To investigate this aAAD paradigm, we designed an experiment where subjects listened to a video lecture under varying attentive conditions. We trained neural decoders to reconstruct the speech envelope from EEG in the baseline attentive condition and use the correlation coefficient between the decoded and real speech envelope as a metric for attention to the speech. Our analysis shows that the envelope standard deviation (SD) of the speech envelope in the 1-4 Hz band strongly correlates with this metric across different segments of the speech stimulus. However, this correlation weakens in the 0.1-4 Hz band, where the degree of separation between the attentive and inattentive state becomes more pronounced. This highlights the unique contribution of the 0.1-1 Hz range, which enhances the distinction of attentional states and remains less affected by confounding factors such as the time-varying dynamic range of the speech envelope.

## I. Introduction

Auditory attention, the ability to focus on a specific speech stimulus, is crucial in classroom and learning settings, where the instructor’s voice often serves as the primary ‘attractor’. However, this attention is not static and fluctuates over time, influenced by factors such as fatigue, distractions, mind wandering, etc. Gaining a deeper understanding of how auditory attention operates and fluctuates in such environments can provide valuable insights into learning processes, including individual differences in how students engage with and process information dynamically.

In the past decade, there has been a lot of research on EEG-based decoding of the speech envelope, in order to decode auditory attention to natural speech. However, most previous studies have focused on *selective* auditory attention (sAAD) decoding, where attention is directed to one of multiple competing stimuli [1]–[5]. While these methods effectively identify which stimulus has been selected, they do not address fluctuations in attention to a single target over time. This gap highlights the need for methodologies capable of *absolute* auditory attention decoding (aAAD)—that is, tracking attention directed solely toward one target stimulus over time.

When capturing fluctuations of attention toward a stimulus, most studies rely on group-level analyses. For instance, metrics like inter-subject correlation (ISC), often computed using Generalized Canonical Correlation Analysis (GCCA), measure synchronization between EEG signals from multiple subjects [6]. While ISC has been shown to correlate with attention, it does not exclusively represent attentional engagement. For example, if subjects are simultaneously distracted by the same external stimulus, their EEG synchronization may remain high despite reduced attention to the relevant target stimulus. To address this issue, a stimulus-aware adaptation of GCCA was proposed in [7]. However, these group-level decoding approaches inherently rely on data from multiple subjects, making them unsuitable for applications focused on individual-level attention tracking.

In the domain of individual attention decoding, metrics such as alpha power [8] and spectral entropy [9] are commonly used. However, these measures are limited as they primarily reflect overall cognitive load rather than attention directed toward a specific (auditory) stimulus [10]. Even when a subject is *not* paying attention to an audio stimulus, their mental effort can still be higher compared to when they are attentively listening [10].

To address these challenges, this work employs individual stimulus-aware aAAD techniques, which use a per-subject neural decoder to reconstruct the speech envelope from EEG, and measure its correlation with the true speech envelope. This correlation coefficient, here referred to as the neural envelope tracking (NET) metric, reflects the strength of the neural response to the speech. This NET metric was shown to be modulated by the degree of attention to the speaker (or distractions from it), for which reason it can be used for aAAD [10].

However, the strength of this neural response could also be affected by other influences rather than attention. One such confound could be the temporal variations in the acoustic dynamic range of the speech signal, reflected by the standard deviation (SD) of the speech envelope. In this paper, we investigate this influence of the envelope SD on aAAD. Our analysis reveals that envelope SD indeed correlates with the NET metric, indicating that temporal variations in the acoustic dynamic range of the speech introduces a confounding effect unrelated to attention. Moreover, this confounding effect varies across different frequency bands of the speech envelope, highlighting the importance of selecting an appropriate band to measure the NET, i.e., the correlation between the real and reconstructed envelope. By selecting the proper frequency band, the NET metric is less affected by external factors such as the time-varying dynamic range in the speech stimulus, allowing it to more accurately reflect genuine cognitive attention.

The paper is organized as follows: Section II introduces the experimental procedures. Section III details the neural decoder design. In Section IV, we present and discuss the experimental results. Finally, Section V concludes the paper.

## II. Data Collection

### A. Subjects

We recruited 30 native Dutch-speaking subjects, including 23 women and 7 men, between 18 and 35 years of age (mean = 24 years; SD = 3 years). None reported any attention deficits. All subjects took part voluntarily and signed an informed consent. All experiments and procedures were approved by the Social and Societal Ethics Committee at KU Leuven.

### B. Protocol Design

The experimental protocol is described in detail in [11]. In summary, we recorded the subjects’ EEG while they were watching a 75-minute course video on the topic of ‘neuromyths’, i.e., widely believed misconceptions about the brain that are actually untrue, based on [12]. It featured a teacher presenting only images in a PowerPoint, with all information delivered orally. The video consisted of 7 parts of approximately 10 minutes, with breaks in between. The experiment included four distinct conditions to explore the dynamics of auditory attention. In the baseline **no manipulation** condition, subjects were instructed to actively listen to the teacher without additional prompts or interruptions. The **dual task** condition required subjects to perform an additional task at the same time, designed to completely divert their attention away from the speaker. These tasks involved solving several calculation exercises, reading a text, or counting down in steps of 17 starting from 994. The remaining two conditions are not discussed as they are beyond the scope of this study and were not analyzed here. The entire timeline is shown in Fig. 1, where each box represents a 30-second window. The white boxes represent the test segments for the ‘no manipulation’ condition, while the red boxes represent the test segments for the ‘dual task’ condition. The gray segments were used for training the decoder (these are all ‘no manipulation’ segments, similar to the white boxes), as explained in Section III. Part 8 is actually a repetition of part 3, but with a continuous dual task during the entire part. The green and purple boxes are other types of manipulations that are not used here.

**Fig. 1:**
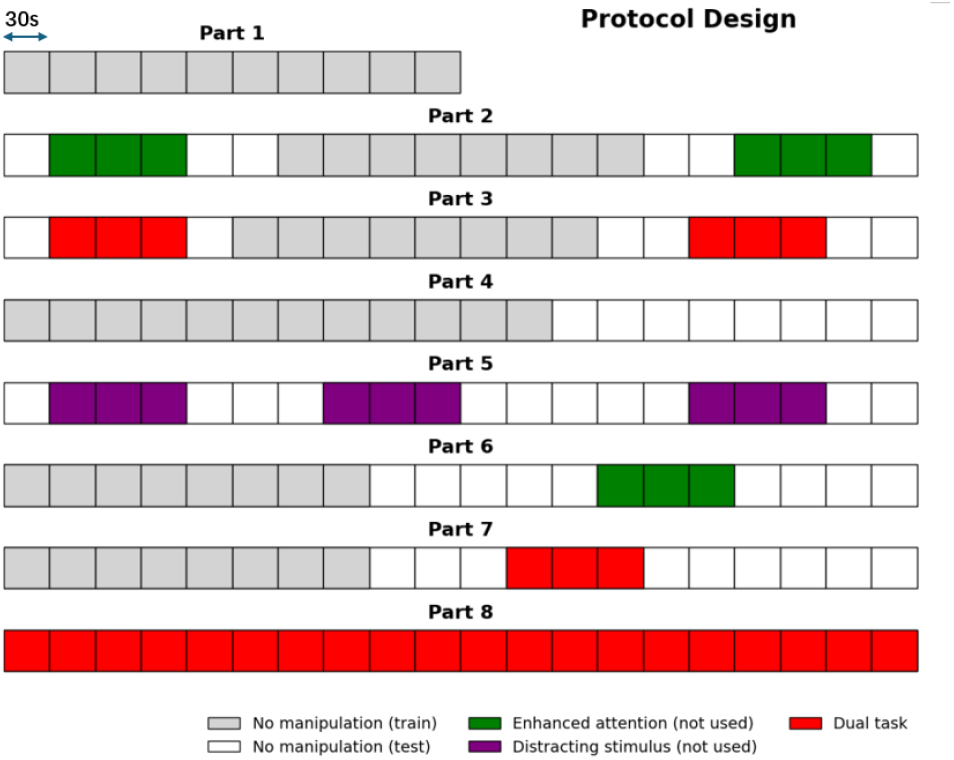
An overview of the protocol. Each box represents a 30-second window.

### C. EEG Data Collection and Preprocessing

The EEG data were collected using a BioSemi Active Two system with 64 electrodes (10-20 system) at a sampling rate of 2048 Hz. The recorded signals were subjected to a series of preprocessing steps to prepare them for analysis.

First, a first-order infinite impulse response (IIR) highpass filter with a cutoff frequency of 0.01 Hz was applied to remove very low-frequency components, such as baseline drift and slow variations. Subsequently, the data was downsampled to 128 Hz to improve computational efficiency while retaining essential EEG features.

Unless specified otherwise, we extracted the 1-4 Hz frequency range proposed in [13], which captures word rates around 2.5 Hz (this is the default setting, we also investigate other choices for the frequency band in Section IV). We used a fourth-order IIR bandpass filter to extract this frequency band, and downsampled the result to a sampling rate of 16 Hz for computational efficiency.

### D. Speech Envelope Extraction

The speech envelope was used as the audio feature in this study, as previous research has shown that certain signal components in the EEG are phase-locked to the speech envelope [1], [9], [10], [13]–[15]. To extract the speech envelope, we used the procedure in [15], which employs a gammatone filterbank to mimic the frequency selectivity of the human cochlea, combined with a power law compression to account for its nonlinear loudness perception.

To align with the EEG preprocessing pipeline, the resulting envelope signal is also bandpass filtered in the same band as the EEG data (1-4 Hz by default) and downsampled to 16Hz.

## III. Neural Decoder Design

Similar to [10], we adopt a spatio-temporal linear decoder that aims to reconstruct the speech envelope of the presented stimulus (i.e., the teacher’s voice) from the EEG of the subject. Formally, we train a decoder 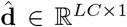 over *T* time samples (corresponding to the training set, i.e., the gray boxes in Fig. 1) such that:

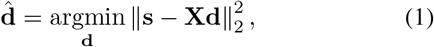

where **s** = [*s*(0) *· · · s*(*T −*1)]^*⊤*^ *∈* ℝ ^*T ×*1^ denotes the *T* samples of the speech envelope, **X** = [**X**_1_ *· · ·* **X**_*C*_] *∈* ℝ ^*T ×LC*^ collects the time-lagged EEG for all *C* channels, and

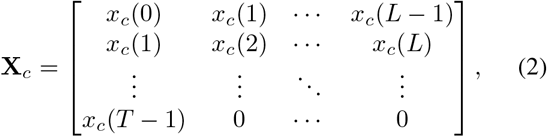

which stacks *L* time-lagged version of the EEG signal *x*_*c*_(*k*) at channel *c*, where *k* is the discrete time (sample) index. A time delay of 0 to 500 ms [13] is chosen corresponding to *L* = 9 (at the 16 Hz sampling rate). Note that this is a noncausal filtering since only post-stimulus time lags are being used. The least-squares solution of (1) is given by

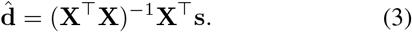

This decoder is trained per subject separately. After training, the decoder is applied to the test segments of the same subject, which is segmented into windows of 30 seconds indexed by *t*. For each window *t*, the EEG data is represented as **X**^(*t*)^ *∈* ℝ ^*N ×LC*^ with *N* = 480 at 16 Hz, and the decoder generates the output 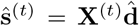. We define the Pearson correlation coefficient between the reconstructed and true stimulus *ρ* (ŝ^(*t*)^, **s**^(*t*)^) as the ‘neural envelope tracking’ (NET) metric to the target speech [10]. It was demonstrated in [10], [13] that higher NET values indicate a higher level of attention to the target speech.

## IV. Analysis and results

### A. Time-varying Envelope SD as a Confound

To investigate the effect of time variations in the dynamic range of the speech stimulus on the NET metric, we computed the standard deviation (SD) of the speech envelope for each 30s segment. Fig. 2a, visualizes the relation between these SD values and our NET attention metric (averaged across subjects) when computed in the default 1-4Hz band from [13], over all test segments (white boxes in Fig. 1). Firstly, as expected, a higher NET metric can be observed for the baseline condition compared to the dual-task condition, which is consistent with [10]. This difference is straightforward to explain: during the dual-task condition, subjects were engaged in another task while ignoring the speech stimulus. This leads to weaker neural tracking compared to the baseline state, resulting in a decrease in the NET metric.

**Fig. 2:**
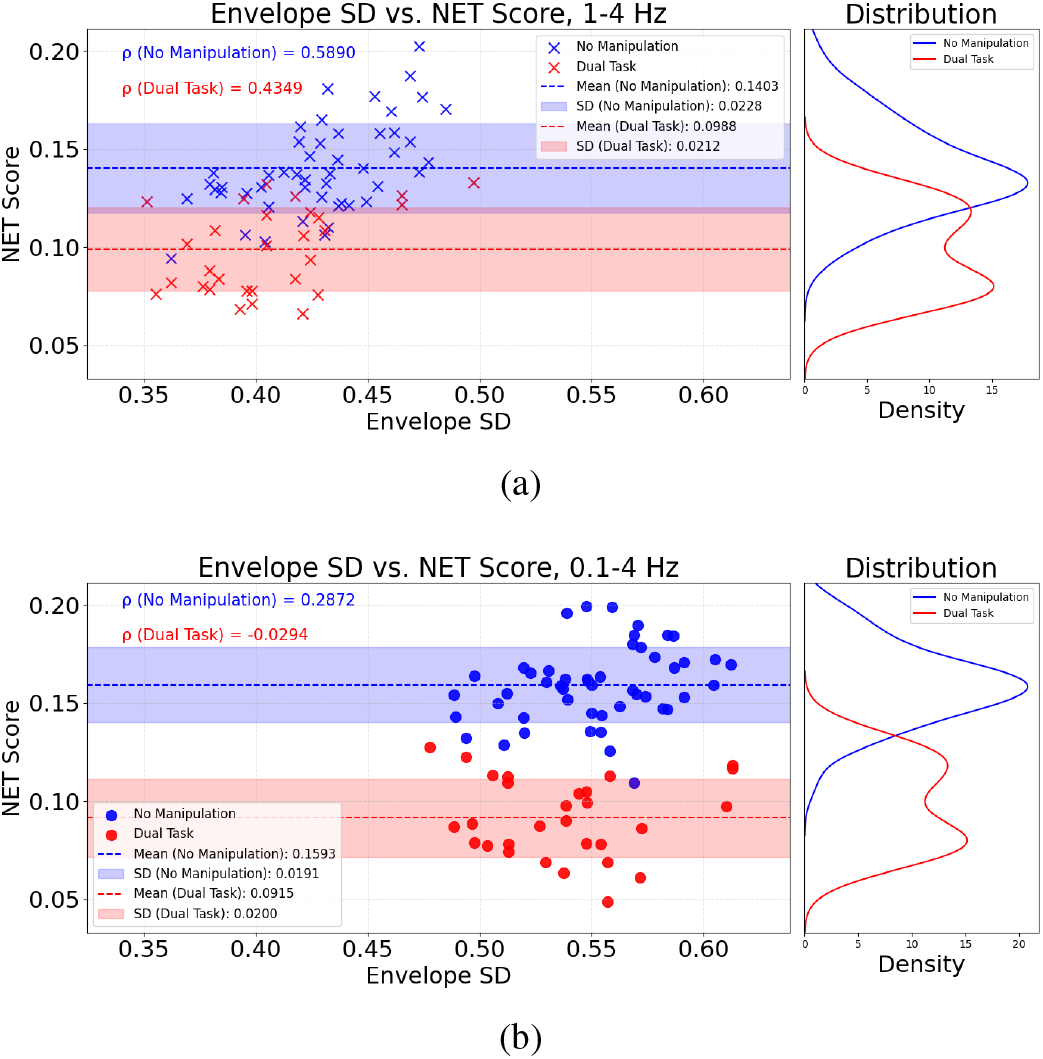
(a) and (b) visualizes the relation between the envelope SD and NET metric for the 1–4 Hz and 0.1–4 Hz frequency bands, respectively. These findings suggest that when attention is focused on the speech stimulus, the influence of envelope SD on neural tracking is more pronounced. Additionally, incorporating the 0.1–1 Hz range reduces this influence while enhancing the ability to distinguish between attention states.

Secondly, we observe that the baseline condition consistently shows stronger correlations between the envelope SD and the NET metric compared to the dual-task condition. For the 1–4 Hz band, the ‘no manipulation’ condition yielded a significant positive correlation (*ρ* = 0.5890, *p <* 0.001), while the ‘dual task’ condition exhibited a weaker correlation (*ρ* = 0.4349, *p* = 0.0184). These results suggest that when attention is focused on the speech stimulus, changes in the dynamic range of the stimulus (represented by the envelope SD) have a stronger effect on the NET. In contrast, the ‘dual task’ condition weakens this relationship as attention is diverted, reducing sensitivity to the envelope SD.

### B. Effect of the Frequency Band on NET

In order to investigate the impact of the frequency band over which the NET is computed (other than the default 1-4Hz from [13]), we have tested different lower and upper cut-off frequencies (between 0.1 and 8Hz), and evaluated the resulting NET metric based on its ability to discriminate between the ‘no manipulation’ and ‘dual task’ conditions. To this end, we use the so-called Fisher Discriminant Ratio (FDR) metric [16], defined as:

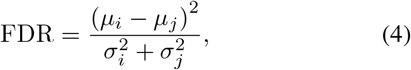

where *i* and *j* represent the two classes (‘no manipulation’ and ‘dual task’), *µ* denotes the mean, and *σ* denotes the standard deviation of the NET scores across all 30-second segments of the corresponding class. A higher FDR value indicates better separability between the baseline and dual task conditions. We found that the band 0.1-4Hz resulted in the highest FDR ratio (averaged across subjects). The relation between the SD envelope and NET metric for this ‘optimized’ frequency band is shown in Fig. 2b.

When comparing Fig. 2a and 2b, we observe that the 1–4 Hz band consistently exhibits higher correlations between the envelope SD and the NET metric than the broader 0.1–4 Hz band for both conditions, particularly in ‘no manipulation’ (*ρ* = 0.5890, *p <* 0.001 for 1–4 Hz; *ρ* = 0.2872, *p* = 0.0454 for 0.1–4 Hz). This highlights the increased sensitivity of the 1–4 Hz range to factors related to the time-varying dynamic range of the speech, which can limit its reliability for decoding attention states. Incorporating the 0.1–1 Hz range in the broader 0.1–4 Hz band reduces this envelope SD correlation values, likely offsetting dynamic range-related effects and making the broader 0.1-4 Hz band more robust for capturing attention fluctuations.

Finally, we observe that the 0.1–4 Hz band better separates the no manipulation and dual task conditions, likely due to reduced sensitivity to dynamic range confounds. As illustrated in Fig. 2, including the 0.1–1 Hz range improves the overall separability of attention states. To quantify this improvement and assess its applicability to individual subjects, we first compute the FDR for each subject and then average these FDR values across subjects. The results show that the mean FDR increases from 0.2195 in the 1–4 Hz band to 0.4484 in the 0.1–4 Hz band, confirming a stronger distinction between attention states. To further analyze the source of this improvement, we examine the mean values of the numerator and denominator in Eq. (4) across subjects. The mean numerator increases substantially (from 0.0027 to 0.0056), while the mean denominator remains nearly unchanged (from 0.0118 to 0.0126). This indicates that the primary factor driving the improvement in FDR is a larger distance between the means of both classes, rather than a reduced intra-class variance. Fig. 3 presents the FDR values for each subject across the two frequency bands. Among the 30 subjects, 25 exhibit an increase in FDR when the 0.1–1 Hz range is included, underscoring the effectiveness of the broader 0.1–4 Hz band for distinguishing attention states.

**Fig. 3:**
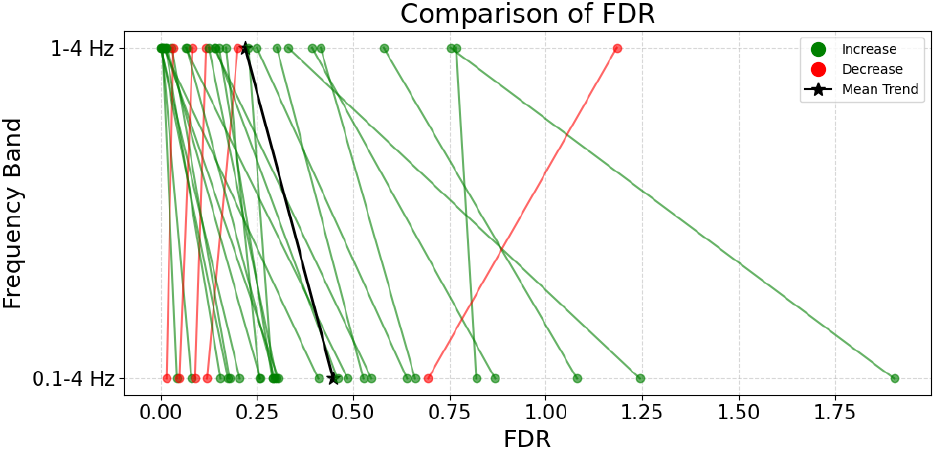
FDR of 1-4 Hz and 0.1-4 Hz frequency bands across subjects. Each dot indicates one subject. Green lines represent an increase, while red lines represent a decrease. The inclusion of the 0.1-1 Hz frequency range leads to an increase in FDR for 25 out of 30 subjects, indicating improved separability of attention states.

## V Conclusion

In this work, we have identified the confounding effect of envelope SD on neural envelope tracking for decoding absolute attention to natural speech. We have then demonstrated that the 0.1-4 Hz components were more robust against this confounding effect compared to 1-4 Hz. Furthermore, we have shown that using the 0.1-4 Hz band improved the separability between attentive and inattentive states.

